# Recombinant chimeric Horsepox Virus (TNX-801) is attenuated relative to Vaccinia Virus Strains in Human Primary Cell Lines and in Immunocompromised Mice

**DOI:** 10.1101/2023.10.25.564033

**Authors:** Stephanie V Trefry, Christy N Raney, Amy L Cregger, Chase A Gonzales, Brittney L Layton, Robert N Enamorado, Nelson A Martinez, Deborah S Gohegan, Tinoush Moulaei, Natasza E Ziółkowska, Scott J Goebel, Seth Lederman, Sina Bavari, Farooq Nasar

## Abstract

Recombinant chimeric Horsepox virus (TNX-801) is a preclinical vaccine in development against Monkeypox and smallpox. In this brief report, we investigated the potential phenotypic differences in *in vitro* and *in vivo* models between TNX-801 and older VACV-based vaccine strains (VACV-IHD, VACV-Lis, VACV-NYC) used in the eradication of smallpox virus. TNX-801 displayed a small plaque phenotype (∼1-2 mm) in both BSC-40 and Vero-E6 cells and yielded >10- to 100-fold lower infectious titers than the VACV strains in multiple-step replication kinetics. Growth kinetics in primary human cell lines from two main routes of poxvirus transmission, respiratory and dermal tracts, yielded ∼10- to 119-fold lower infectious titers of TNX-801. Intranasal infection of immunocompromised mice (C56BL/6 *Ifnar*^−/−^/*Ifngr*^−/−^) with VACV strains at 6.0 and 5.0 log_10_ PFU produced uniform lethal disease. In contrast, TNX-801 at 8.0 log_10_ PFU was unable to produce any clinical disease in mice. These data demonstrate that TNX-801 is >10- to 1,000-fold more attenuated than older VACV-based smallpox vaccines in human primary cell lines and immunocompromised mice.

## Introduction

Horsepox virus (HPXV) is a member of the genus *Orthopoxvirus* in the family *Poxviridae*. The genome is comprised of double-stranded DNA ∼212 kb in length [1]. Little is known about the biology of HPXV; however, the genus is also home to four prominent members, Variola, monkeypox, cowpox, and vaccinia viruses, which are important in human health. Variola, the causative agent of smallpox disease, and monkeypox viruses are important human pathogens that can cause fatal infection. Case fatality rates with smallpox and monkeypox infection range from ∼30% to ∼10%, respectively [1]. Following a world-wide vaccination campaign, smallpox was eradicated in 1980. The smallpox vaccine was the product of Edward Jenner’s seminal observation in milkmaids who contracted a mild disease from the cows and in turn were protected from lethal smallpox. For 130 years following his observation, the virus in the vaccination efforts was thought to be of Cowpox origin [1-3]. However, in 1939, it was shown that the virus in the vaccine was comprised of Vaccinia Virus (VACV), a virus serological related but distinct from Cowpox. The origin of VACV, its natural cycle, and how it was introduced remain unknown. Recent genomic studies of American Civil War Era smallpox vaccine scab kits showed the presence of both VACV and HPXV [4]. One of the isolates displayed a 99.7% nucleotide identity to HPXV [4]. These data strongly support Jenner’s hypothesis and suggest that HPXV was used as a human vaccine. Utilizing modern genetic techniques, we generated an infectious clone of Horsepox virus (TNX-801) and are investigating it as a next generation vaccine against smallpox and monkeypox disease [5]. One critical hurdle in the development of any live-attenuated vaccine platform is ensuring safety in relevant model systems. Preliminary studies demonstrated that TNX-801 readily grew in mammalian cells, however, it displayed an attenuated phenotype both *in vitro* and *in vivo* models [5]. In this study, we aimed to extend the previous findings by investigating the attenuation of TNX-801 in immortalized non-human primate (NHP) cell lines, primary human cells, and in an immunocompromised mouse model relative to the older Vaccinia virus-based smallpox vaccines (VACV-IHD, VACV-Lister, and VACV-NYC) utilized to eradicate smallpox.

## Results

### TNX-801 plaque phenotype

The plaque phenotype of TNX-801 was compared relative to VACV-WR, VACV-IHD, VACV-Lis, VACV-NYC, and MVA on immortalized non-human primate cell lines, BSC-40 and Vero-E6 cells (Figs. 1 and 2). In both cell lines, VACV strains displayed large plaque phenotype plaques ∼3-4 mm in diameter by 4 (BSC-40) and 6 dpi (Vero-E6). In contrast, TNX-801 and MVA displayed small plaque phenotype with ∼1-2 mm and ∼0.5 mm in diameter on both cell lines, respectively.

**Figure 1.**
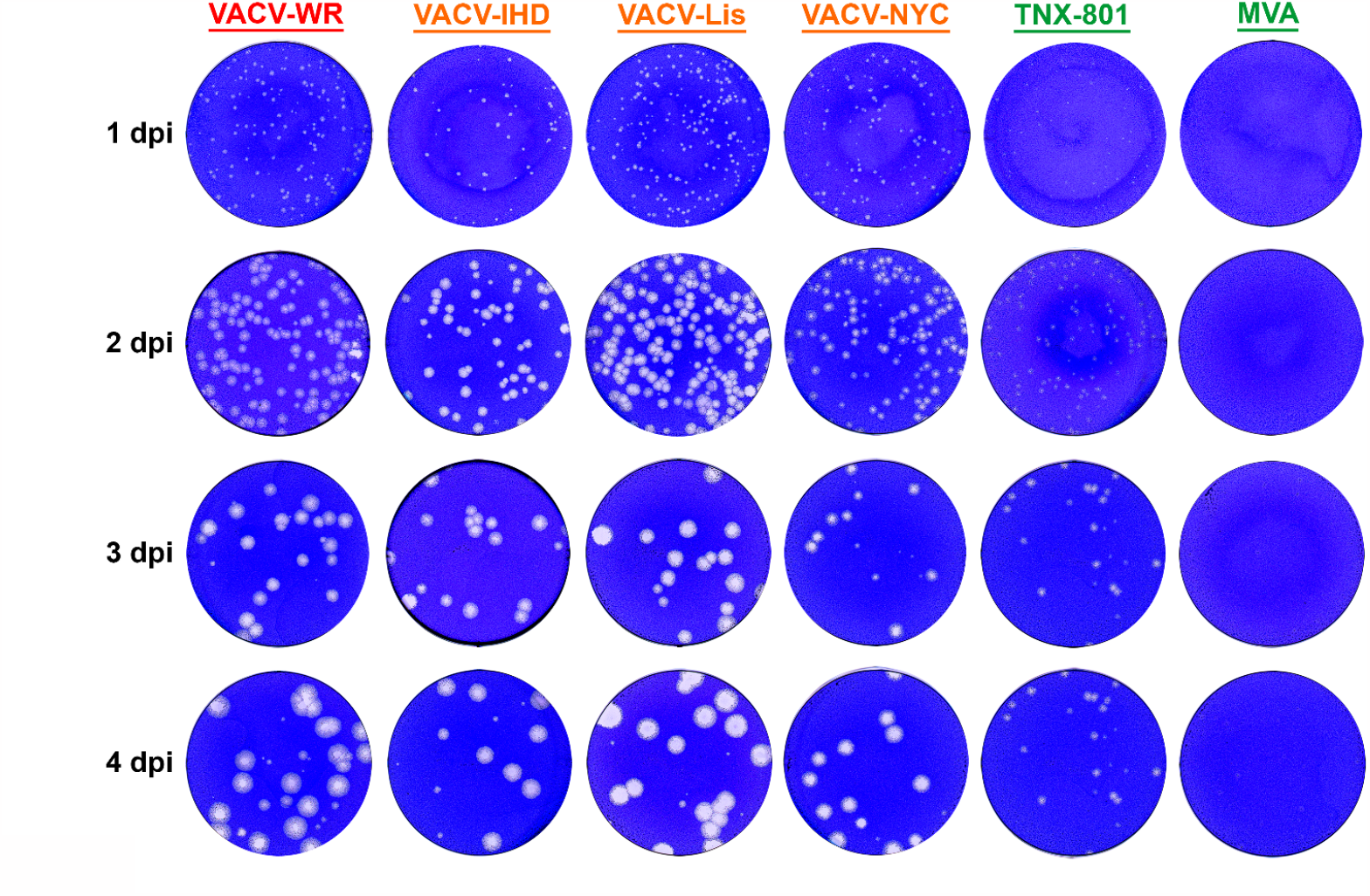
Plaque phenotype of VACV-WR, VACV-IHD, VACV-Lis, VACV-NYC, TNX-801, and MVA was investigated in BSC-40 cells. Representative wells from 6-well plates are shown in each photograph.

**Figure 2.**
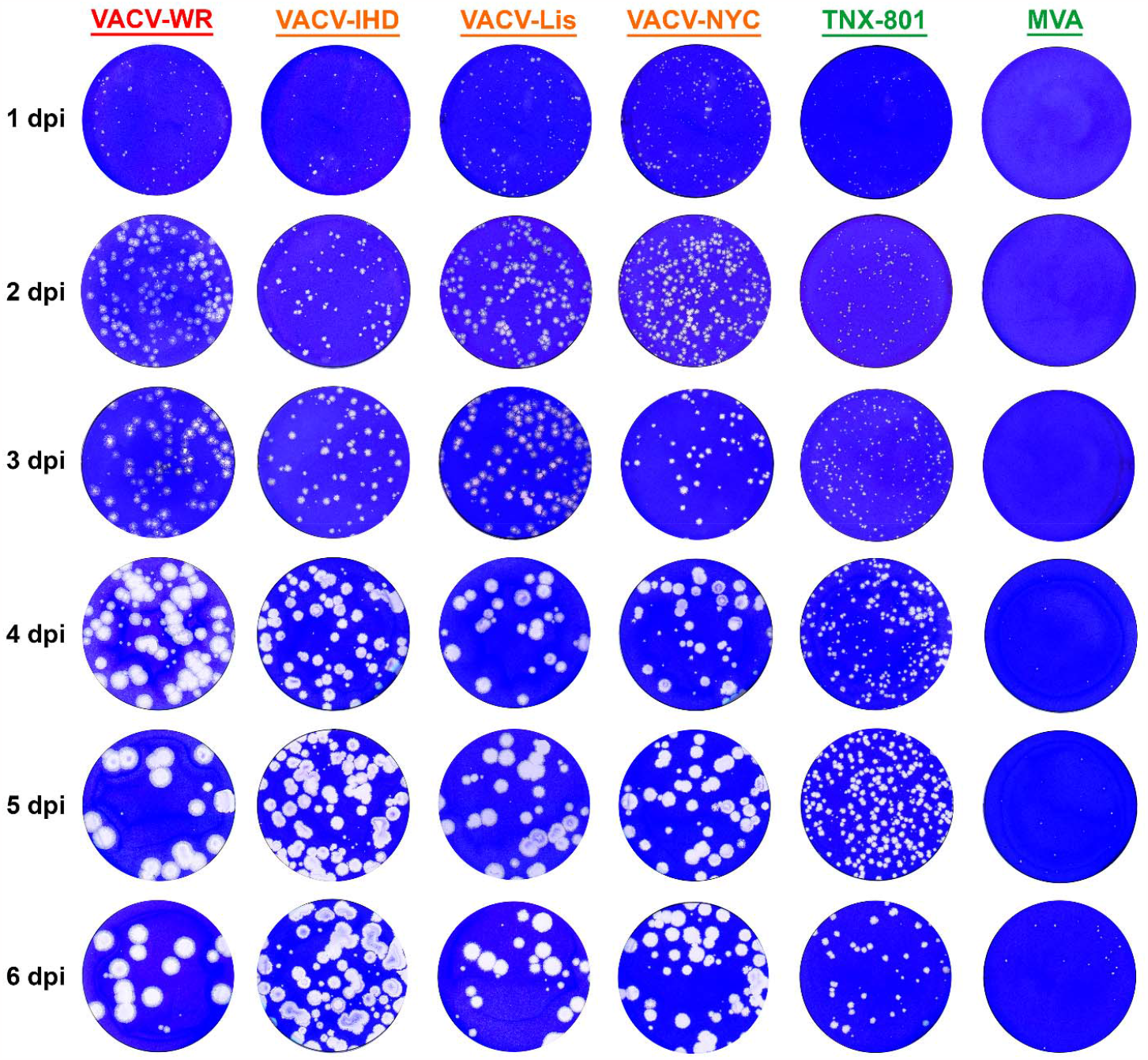
Plaque phenotype of VACV-WR, VACV-IHD, VACV-Lis, VACV-NYC, TNX-801, and MVA was investigated in Vero-E6 cells. Representative wells from 6-well plates are shown in each photograph.

### Multiple-step replication kinetics

The replication kinetics of TNX-801, VACV-WR, VACV-IHD, VACV-Lis, VACV-NYC, and MVA was investigated in BSC-40 and Vero-E6 cells at a multiplicity of infection (MOI) of 0.01 (Figure 3A and B). All VACV strains reached peak titers of ∼8.0 log_10_ PFU/mL between 48 and 72 hpi. In contrast, TNX-801 and MVA displayed delayed replication kinetics reaching peak titers between 72 and 96 hpi. The peak titer of TNX-801 was ∼7.7 and ∼6.6 log_10_ PFU/mL in BSC-40 and Vero-E6 cells, respectively. The TNX-801 titers were up to ∼10-to 119-fold lower than VACV strains timepoints. MVA infection yielded peak titers of ∼5.6 and ∼4.8 log_10_ PFU/mL in BSC-40 and Vero-E6 cells, respectively.

**Figure 3.**
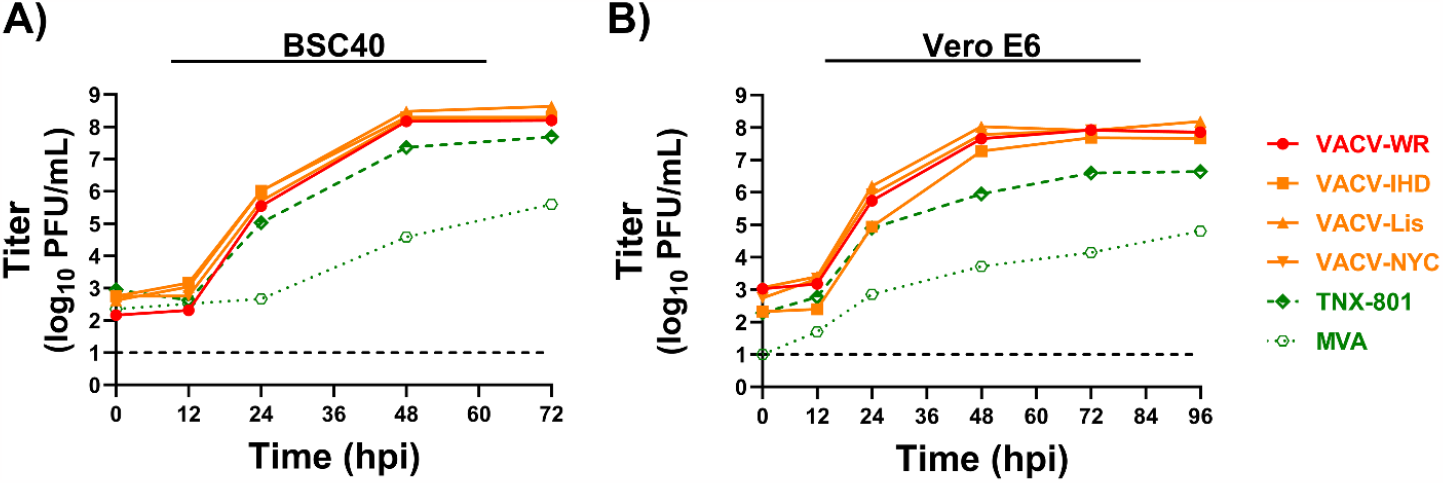
Multi-step growth kinetics of VACV-WR, VACV-IHD, VACV-Lis, VACV-NYC, TNX-801, and MVA in BSC-40 (A) and Vero-E6 (B) cells. The limit of detection is shown with black dashed line.

Multiple-step replication studies were also performed in human primary cell lines representing two main routes for poxvirus transmission from the dermal (melanocytes, keratinocytes, dermal fibroblasts, and skeletal muscle) and respiratory (bronchial/tracheal epithelial, small airway epithelial, lung fibroblasts, and bronchial/tracheal smooth muscle) tracts. Cells were infected at an MOI of 0.01 and samples were collected at 24-hour (hrs) intervals (Fig. 4A). The TNX-801 infection of primary melanocytes, keratinocytes, and fibroblasts displayed delayed replication kinetics. The peak infectious titer of TNX-801 and VACV strains ranged from ∼3.8 to 5.9 in log_10_ PFU/mL and ∼4.5 and 7.5 log_10_ PFU/mL, respectively. The infectious titers of TNX-801 were ∼5-to 112-fold lower in all three primary cell lines. In contrast, TNX-801 and VACV strains yielded similar infection kinetics and infectious titers in skeletal muscle cells with peak titers of ∼7.1 and ∼7.7 log_10_ PFU/mL. The MVA infection did not produce any infectious virus in any dermal tract primary cells throughout the 96-hr sampling period.

**Figure 4.**
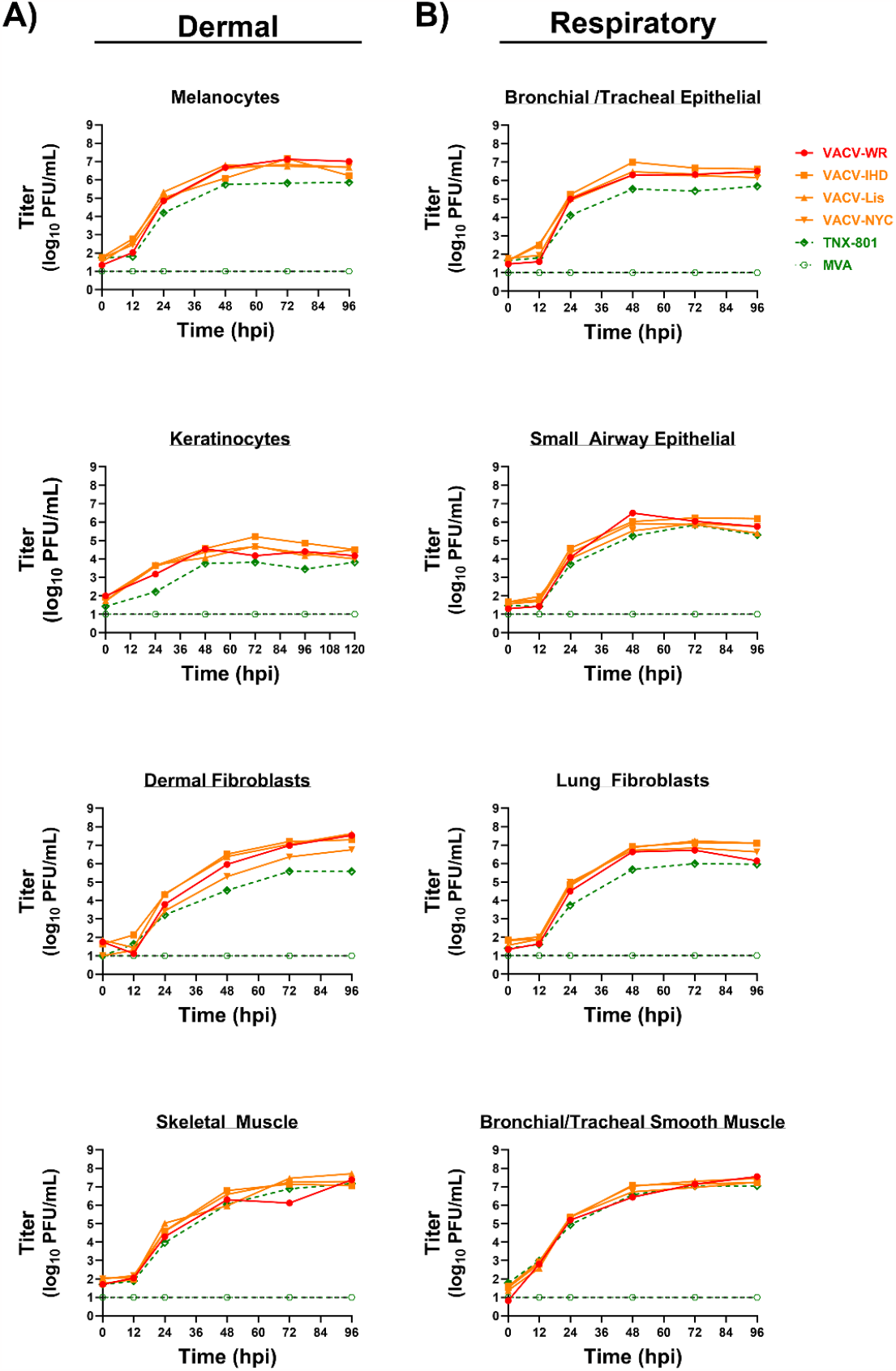
Multi-step growth kinetics of VACV-WR, VACV-IHD, VACV-Lis, VACV-NYC, TNX-801, and MVA in primary human cell lines from dermal (A) and respiratory (B) tracts. The limit of detection is shown with black dashed line.

Infection of primary cells from the respiratory tract yielded similar results. The TNX-801 infection of fibroblasts, bronchial tracheal epithelial and small airway epithelial cells produced peak infectious titers ranging from ∼5.6 to 6.0 log_10_ PFU/mL (Fig 4B). VACV strains yielded peak infectious titers of ∼6.0 to 7.2 log_10_ PFU/mL. TNX-801 titers were ∼5-to 28-fold lower than VACV strains. In contrast, TNX-801 and VACV strains displayed similar infection kinetics and infectious titers in bronchial tracheal smooth muscle cells with peak titers of ∼7.0 to 7.5 log_10_ PFU/mL. Lastly, infection with MVA did not produce any infectious virus in any respiratory tract primary cells throughout the 96 hr sampling period.

### Intranasal infection studies in C56BL/6 Ifnar^−/−^/Ifngr^−/−^ mice

To further investigate phenotypic differences between TNX-801 and VACV strains, susceptibility to virus infection was determined in C56BL/6 *Ifnar*^−/−^/*Ifngr*^−/−^ mice via the intranasal route (Fig 5A). Cohorts of 10 mice/group were infected at 6.0 and 5.0 log_10_ PFU/mouse with VACV-WR, VACV-IHD, VACV-Lis, and VACV-NYC. Mice were also infected with TNX-801 at 8.0 log_10_ PFU/mouse or mock infected with phosphate buffered saline (PBS) (Fig 5A).

**Figure 5.**
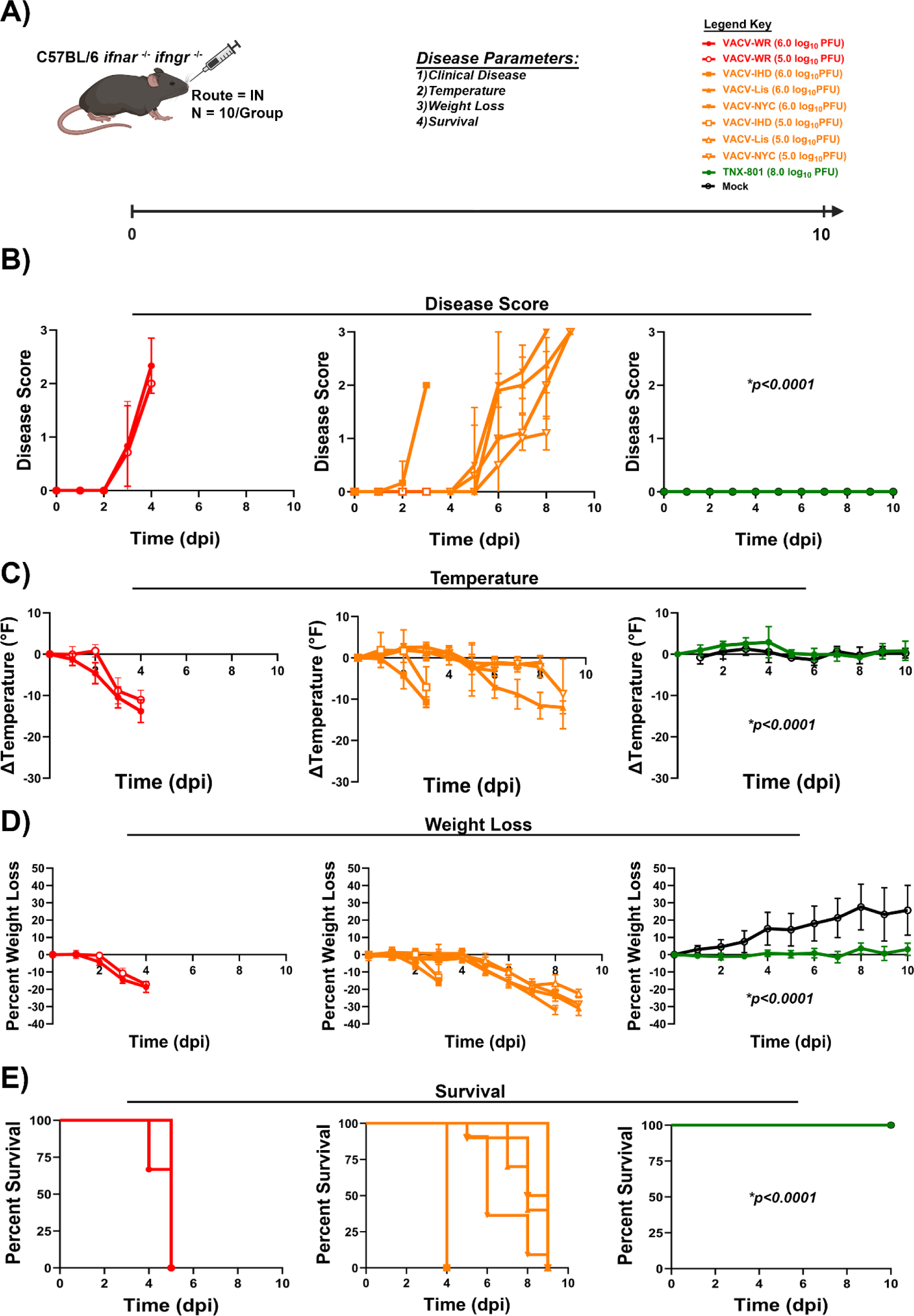
Infection of C57BL/6 *ifnar* ^−/−^/*ifngr* ^−/−^ mice with VACV-WR, VACV-IHD, VACV-Lis, VACV-NYC, TNX-801, and mock treated via intranasal route (A). Mice were infected with VACV-WR, VACV-IHD, VACV-Lis, and VACV-NYC at 6.0 and 5.0 log_10_ PFU/mouse. Mice were infected with TNX-801 at 8.0 log_10_ PFU/mouse. Post-infection, mice were monitored for clinic disease (B), body temperature (C), weight loss (D), and survival (E). *p*-values comparing TNX-801 to VACV strains are provided in each graph.

Following infection with VACV-WR, VACV-IHD, VACV-Lis, and VACV-NYC, VACV-WR and VACV-IHD infected mice exhibited accelerated clinical disease. All mice exhibited clinical disease by 3 days post-infection (dpi) with peak scores of 2.3 and 2.0 for VACV-WR and VACV-IHD, respectively (Fig 5B). Body temperature began to decline between 1 to 2 dpi then accelerated with average peak decline ranging from −11°F to −14°F at 3 to 4 dpi (Fig 5C). Similar kinetics were observed for weight loss with peak average weight loss of ∼17% (Fig 5D). All mice met the euthanasia criteria between 4 and 5 dpi regardless of dose (Fig 5E). Mice infected with VACV-Lis and VACV-NYC exhibited delayed disease kinetics. The onset of clinical disease was between 5 and 6 dpi, and the peak average scores ranged from 2.2 to 3 (Fig 5B). Body temperature declined between 5 and 6 dpi with peak average decline ranging from −2°F to −12°F (Fig 5C). Similarly, weight loss began to decline at 5 dpi and the peak average weight loss ranged from ∼22% to 32% (Fig 5D). All mice met the euthanasia criteria between 5 and 10 dpi (Fig 5E). In contrast to all VACV infected mice, TNX-801 and mock infected mice did not exhibit any clinical disease or change in body temperature and weight loss, and all mice survived the study (*p<0*.*0001*).

## Discussion

Recombinant chimeric HPXV (TNX-801) vaccine is a preclinical stage vaccine against monkeypox and smallpox infection. One critical hurdle in the development of replication competent vaccines is demonstration of vaccine safety in relevant *in vitro* and *in vivo* models. In this study, we investigated the potential attenuation of TNX-801 relative to VACV-based smallpox vaccines used to eradicate smallpox. In immortalized non-human primate, TNX-801 displayed a small plaque phenotype, delayed replication kinetics, and >10-to 100-fold lower infectious titers. Human primary cells from dermal and respiratory tracts also showed similar levels of attenuation. Lastly, in the infection of immunocompromised mice, TNX-801 was unable to cause any clinical disease, alteration in body temperature and weight, and all mice survived the study period. Taken together, these data demonstrate that TNX-801 is 10-to 1,000-fold more attenuated than older Vaccinia virus-based vaccines used in smallpox eradication.

TNX-801 displayed an attenuated phenotype in majority of the human primary cells from two main poxvirus transmission routes, dermal and respiratory tracts. However, in skeletal muscle and bronchial tracheal smooth muscle cells TNX-801 displayed comparable fitness to VACV. These data suggest that the TNX-801 attenuation is more prominent in cells that would serve as initial site of replication during virus infection and will limit the spread of TNX-801 by either dermal or respiratory route. The lack of clinical disease in mice supports this hypothesis. Surprisingly, TNX-801 displayed comparable replication kinetic and infectious titers in primary human skeletal muscle cells. This suggest that the intramuscular route may provide superior immunogenicity following vaccination with TNX-801. Vaccination studies in rabbits and NHPs with TNX-801 has focused on scarification route and have yielded robust humoral responses [6]. Future studies will investigate multiple vaccination routes and determine the optimal route of vaccination in human clinical trials.

Interferon alpha and gamma play a critical role in host response and defense against viral pathogens. The disruption of either or both pathways enable susceptibility in mice to virus infection. We investigated an immunocompromised mouse model comprising of interferon alpha and gamma receptor knockout to enable increased susceptibility to poxviruses. The mouse model was extremely susceptible to infection with VACV strains with all mice rapidly succumbing to virus infection at both 6.0 and 5.0 log_10_ PFU doses. In contrast, mice infected with TNX-801 at a 1,000-fold higher dose were unable to exhibit any clinical disease. These data strongly suggests that the vaccine platform is significantly more attenuated than VACV strains. Studies are underway to determine the lethal dose (LD_50_) of all VACV strains. In addition, future studies will explore virus dissemination and virus induced pathology.

The central core of poxvirus genome ∼90 kb in length encodes genes responsible for virus replication and consequently is highly conserved [1, 7]. However, the left and right arms of the genome are more flexible and encode for host range restriction and/or host immune antagonism genes. TNX-801 genome is ∼20 kb larger than VACV genome, and yet it displays an attenuated phenotype than VACV [1, 7]. The gene/s that underlie this phenotype are under investigation and future studies will aim to determine their effect on various stages of poxvirus replication cycle.

In conclusion, the potential phenotypic difference of TNX-801 and VACV strains was investigated in *in vitro* and *in vivo* models. TNX-801 displayed an attenuated phenotype relative to VACV with a small plaque morphology and >∼10-to 100-fold lower infectious titers in immortalized and primary cell models. In immunocompromised mice, TNX-801 was >1,000-fold more attenuated than VACV-based vaccines utilized for the eradication of smallpox.

## Materials and Methods

### Cells

Vero (CRL-1586) and BSC-40 (CRL-2761) cells were obtained from American Tissue Culture Collection (ATCC, Manassas, VA). Cells were cultured in Dulbecco’s MEM (DMEM) supplemented with 10% fetal bovine serum (FBS) and gentamicin (50 μg/mL) at 37°C and 5% CO_2_.

Normal adult human primary cells were obtained from American Tissue Culture Collection (ATCC, Manassas, VA). Cells selected were representative of two transmission routes, respiratory or dermal tracts. Primary dermal tracts cells comprised of Epidermal Melanocytes (HEMa, PCS-200-013), Epidermal Keratinocytes (HEKa, PCS-200-011), Dermal Fibroblasts (HDFa, PCS-201-012) and Skeletal Muscle (HSkMC, PCS-950-010). Primary respiratory tract cells were Bronchial/Tracheal Epithelial (PCS-300-010), Small Airway Epithelial (HSAEC, PCS-301-010), Lung Fibroblasts (HLF, PCS-201-013), and Bronchial/Tracheal Smooth Muscle (PCS-130-011). Cells were cultured in their respective basal mediums and growth kit supplements per manufacturer recommendations. All primary cells were cultured at 37°C and 5% CO_2_. Cells were seeded overnight to achieve appropriate confluence for the multi-step growth kinetics.

### Viruses and Virus Amplification

Vaccina virus strains [Lister (Lis), International Health Department (IHD), Western Reserve (WR), New York City (NYC), and Modified Vaccinia Ankara (MVA) were obtained from BEI Resources (Manassas, VA). HPXV (TNX-801) stock rescued as described previously. Virus stocks of each strain were generated on BSC-40 cells. Briefly, cells were seeded overnight in flasks to achieve 70% confluence. Growth media was removed, and monolayers were infected at an MOI of 0.01. Following an hour incubation at 37°C and 5% CO_2_, 20 mL growth media was added. Monolayers were harvested via cell scraping 72 hpi. Samples were subjected to three cycles of freeze/thaw and sonication and were stored at −80°C. All virus stocks were titrated by plaque assay on BSC-40 cells.

For mouse studies, all virus stocks were sucrose purified. Briefly, SW-28 tubes were filled with 16 mL of 36% sucrose (w/v) in 10mM Tris pH 7.5 followed by overlay of 16 mL of virus isolates. Tubes were spun at 4°C at 33,000 x *g* for 80 mins. Following centrifugation, all supernatants were discarded, and the virus pellet was resuspended in 5 mL of 1X PBS. Sucrose purified virus was titrated on BSC-40 cells.

### Virus Titration

BSC-40 cells were seeded overnight in 6-well plates to achieve 95-100% confluence. Each virus stock was serially diluted 10-fold in 1X DMEM. Monolayers were infected with 100 μL of each dilution and incubated at 37°C and 5% CO_2_ for one hour. Following incubation, monolayers were overlaid with 2 mL of 0.5% methylcellulose in MEM, GlutaMAX™ (Gibco) supplemented with 5% FBS and gentamicin (50 μg/mL). Following 72 hours at 37°C and 5% CO_2_, plates were fixed in 10% formalin overnight. Plates were stained with 2% crystal violet in 70% ethanol for 5-10 minutes. Stain was removed with water, plates were dried, and plaques were counted. For plaque phenotype comparison, the identical protocol was performed on BSC-40 and Vero-E6 cells and representative 6-wells are shown in figures.

### Multi-Step Growth Kinetics

BSC-40, Vero-E6, and human primary cells were seeded overnight in 6-well plates to achieve 50% confluence. Growth media was removed, and monolayers were infected at an MOI of 0.01 with TNX-801, VACV-WR, VACV-IHD, VACV-Lis, VACV-NYC, and MVA. Following an hour incubation at 37°C and 5% CO_2_, monolayers were washed with 10 mL PBS, and 2 mL of growth media was added. Monolayers were harvested via cell scraping at 0, 12-(BSC-40 and Vero-E6 cells only), 24-, 48-, 72-, and 96-hpi in triplicate. Samples were subjected to three cycles of freeze/thaw and sonication, and were stored at −80°C. All samples were titrated on BSC-40 cells.

### Ethics statement

This work was supported by an approved Tonix Pharma Animal Care and Use Committee (IACUC) animal research protocol. Research conducted under an IACUC approved protocol and was in compliance with the Animal Welfare Act, PHS Policy, and other Federal statutes and regulations relating to animals and experiments involving animals. The facility where this research was conducted is accredited by the Association for Assessment and Accreditation of Laboratory Animal Care (AAALAC International) and adheres to principles stated in the Guide for the Care and Use of Laboratory Animals, National Research Council, 2011 [8].

### Mouse Studies

Breeding pairs of C57BL/6 *ifnar* ^−/−^/*ifngr* ^−/−^ were purchased from Jackson Laboratories to generate mouse colonies at Tonix Pharma. A ten-day study was performed to investigate virulence of TNX-801, VACV-WR, VACV-IHD, VACV-Lis, and VACV-NYC. Cohorts of ten (5 male and 5 female) four-week-old mice were infected via intranasal route. Mice were infected with 50 μL (25 μL/nare) of virus. Mock infected mice received 50 μL (25 μL/nare) of 1X PBS. C57BL/6 *ifnar* ^−/−^ /*ifngr* ^−/−^ mice were infected with VACV-WR, VACV-IHD, VACV-Lis, and VACV-NYC at 6.0, and 5.0 log_10_ PFU/mouse. Mice were infected with TNX-801 at 8.0 log_10_ PFU/mouse. Following infection, mice were monitored daily for clinical disease, body temperature, weight loss, and survival. The clinical disease score criteria: 0 = normal, 1 = rough coat and/or hunched posture, 20% body weight loss from baseline, 2 = mild lethargy and/or mild dyspnea, 25% body weight loss from baseline, and 3 = moderate lethargy and/or moderate dyspnea and/or 30% body weight loss from baseline. All target doses were verified via plaque assay and actual dose is shown in respective figures.

## Statistical Analysis

GraphPad Prism version 9 for Windows (GraphPad Software, La Jolla, California, USA) software was utilized for statistical analysis. Significant differences in each parameter were determined using one-way ANOVA followed by multiple comparisons between groups. Significance between TNX-801 and VACV strains were reported in the graphs.

## Disclaimer

TNX-801 vaccine is not approved for any indication.

